# N-terminal helices and A domain of archaeal FtsY facilitate SRP54 binding and the association with cell membrane

**DOI:** 10.1101/2022.04.13.488184

**Authors:** Sayandeep Gupta, Souvik Sinha, Koustav Bhakta, Arghya Bhowmick, Abhrajyoti Ghosh

## Abstract

The process of protein translocation is essential to the maintenance of cellular life and has been critically addressed in eukaryotes and bacteria. However, little information is available regarding protein translocation across archaeal membranes. The signal recognition particle (SRP) plays an important role in this process. It binds the signal peptide at the N-terminus of the polypeptide chain and interacts with the cognate SRP receptor (FtsY) located on the target membrane to form a targeting complex (TC). Concomitant GTP hydrolysis by SRP and FtsY delivers the polypeptide to the adjacent protein-conducting channel. The present study aims to characterize the structural domains of FtsY contributing to the targeting complex (TC) formation in *Sulfolobus acidocaldarius*, a thermo-acidophilic crenarchaeon. The contacting residues between SRP54 and FtsY were mapped along the αN1-N3 helices. Interestingly, the previously reported crystal structure did not take the N-terminal A domain into account – a region rich in negatively charged residues. Such observation led us to investigate the contribution of each of the three participating helices (αN1-3) in terms of membrane association and functional TC formation. Through biophysical analyses of SRP-FtsY and FtsY-membrane interaction, and biochemical characterization of the reciprocal GTPase activity, this work sought to elucidate the minimal structural motif controlling the archaeal TC assembly.

## 1. Introduction

Archaeal SRP receptor shares structural similarity with the bacterial counterpart, FtsY, which itself is a homolog of the alpha subunit of eukaryotic SRP receptor (SIRα) (Miller et al., 1994). It has an N-terminal A domain consisting of acidic (negatively charged) residues, followed by a conserved NG domain, the same as the one shared by SRP54 (Gupta et al., 2016). The classical GTPase motifs (I-V) are also shared by the two SRP GTPases, along with the IBD and other signature motifs (Keenan et al., 2001; Leipe et al., 2002) but contrary to the classical P-loop GTPases, they do not employ a guanine nucleotide exchange factor and directly interact with each other to stimulate the concomitant hydrolysis of GTP (Powers & Walter, 1995; Jagath et al., 2000). Membrane targeting by the RNC-SRP complex is facilitated via multiple conformational changes brought about by the dynamic interaction between NG domains of the SRP GTPases (Shan et al., 2007; Zhang et al., 2009). This interaction was found to be strengthened by the helix 8 tetraloop of the SRP RNA where the SRPoGTP-GTPoSR complex initially assembles and then relocalizes to the distal end where the GTP hydrolysis is stimulated (Ataide et al., 2011; Shen et al., 2012; Shen et al., 2013). The “twinned” hydrolyses of the bound GTPs promote the dissociation of the Ffh-FtsY targeting complex and thus the cargo is unloaded upon the translocon (Egea et al., 2004; Shan et al., 2004).

Structural analyses of the Ffh-FtsY complex have established the contribution of NG domains of the two proteins in their mutual interaction. The heterodimeric complex from *Thermus aquaticus* revealed that the primary interactive interface exists between the G domains, accompanied by the interaction between N domains, comprising extensively of hydrogen bonds and van der Waals contacts (Egea et al., 2004). Individual mutagenesis of the residues along the interaction surface of FtsY in *E. coli* severely depleted the reciprocally stimulated GTP hydrolysis by Ffh-FtsY but the basal GTPase activity of FtsY was unaffected (Egea et al., 2004). The association of the targeting complex and its dual GTPase activity depended extensively on the full-length SRP RNA both in bacteria and in archaea (Zhang et al., 2008; Gupta et al., 2021). Crystal structure of the bacterial TC showed profound domain rearrangement of Ffh upon binding the 4.5S RNA at the tetraloop region via the M domain leading to the movement of the NG domain to the distal region of helix 5 (Ataide et al., 2011). This was further substantiated by the binding analyses between NG and M domains of Ffh and FtsY in the presence or absence of SRP RNA (Buskiewicz et al., 2005). The binding of FtsY-NG with the SRP54/Ffh-NG is facilitated due to this positional reorientation in the latter and enhanced greatly by the presence of GTP (Jagath et al., 2000; Gupta et al., 2021). Together, the two reoriented NG domains form a composite active center of the TC that binds and hydrolyses GTP, and finally, the activated TC complexed with the RNC interacts with the membrane-bound translocon SecYEG via FtsY (Draycheva et al., 2016).

Although the Ffh-associating NG domain of FtsY is quite conserved, the unstructured A domain varies among different homologs of the prokaryotic SRP receptor (Bibi et al., 2001). The eukaryotic homolog SIRα, which carries out similar functions as FtsY, remains associated with its integral membrane-bound component SRβ (Walter & Johnson, 1994). The absence of such a membrane-spanning counterpart and the evidence of FtsY being distributed between the cytoplasm and the membrane have confirmed the soluble nature of the receptor in bacteria (Luirink et al., 1994). Therefore, it was suggested that the A domain may have a role in membrane association (Powers & Walter, 1997; Zelazny et al., 1997), though both A and NG domains of bacterial FtsY have been shown to have an affinity toward membrane phospholipid (de Leeuw et al., 2000). However, the A domain in *E. coli* was shown to be necessary for targeting the NG domain to the membrane and carrying out efficient translocation (Powers & Walter, 1997). Functional replacement of this domain with an unrelated integral membrane polypeptide proved the importance of the acidic domain in membrane association (Zelazny et al., 1997). Surprisingly, deletion of the A domain (1-195 residues) from *Ecoli*FtsY, termed NG+1, did not affect its cellular functions significantly, but deletion of a single residue (Phe196) lead to a completely inactive NG construct (Bahari et al., 2007; Draycheva et al., 2016). FtsY-NG retained its *in vitro* basal GTPase and Ffh-binding activities and accumulated in the plasma membrane along with SRP-RNC, indicating that the release of the complex from the membrane was indeed defective (Bahari et al., 2007). This led the researchers to believe that the interaction with lipid is not essential for either membrane-targeting or receptor-docking, but rather for efficient release of the complex upon cargo unloading. This was further confirmed by the finding that anionic phospholipid induced conformational changes in FtsY which stimulated its basal GTPase activity (de Leeuw et al., 2000; Lam et al., 2010) and facilitated the formation of the GTP dependent intermediate of TC (Lam et al., 2010). The N-terminal alpha-helices in both proteins play major roles in complex assembly and function, in fact, deletion of αN1 of FtsY greatly enhanced the complex formation and the basal GTPase activity as seen in the presence of phospholipid (Stjepanovic et al., 2011). Thus, phospholipid may induce the conformational change in the αN1 helix that is part of the proposed membrane targeting sequence (MTS).

In the haloarchaeal SRP system, the *in vivo* functionality of the FtsY was found to be similar both in the presence and absence of the A domain (Haddad et al., 2005). Structural studies involving the FtsY from *Pyrococcus furiosus* (Egea et al., 2008) and *Sulfolobus acidocaldarius* (Wild et al., 2016) have contributed considerably to the otherwise scarce pool of knowledge about archaeal SRP receptors. The *PfuFtsY* showed an elongated αN1 helix instead of the unfolded A domain commonly found in bacteria whereas *SaciFtsY* had a shorter αN1 and the contacting residues between SRP54 and FtsY were mapped along the αN1-N3 helices. Interestingly, the crystal structure did not employ the initial 20 amino acids at the N-terminus into account -a region rich in negatively charged residues. Such observation led us to investigate the contribution of each of the three participating helices (αN1-3) in terms of membrane association and functional TC formation. The lack of proper biochemical evidence and an in-depth understanding of the minimal functional MTS and its mechanism of membrane targeting in archaea have been the major driving factor for this work. By utilizing a vast array of biophysical techniques such as fluorescence resonance energy transfer, circular dichroism, anisotropy, and biochemical characterization of the reciprocal GTPase activity, we sought to elucidate the protein-protein and lipid-protein interactions controlling the archaeal TC assembly.

## 2. Materials & methods

### 2.1. Construction of overexpression vectors

The construction of pAG3 and pAG351, expressing wild type SRP54 and FtsY respectively, was described earlier (Gupta et al., 2021). The deletion mutants were generated using the pAG351 backbone following site-directed mutagenesis protocol. Primers used for this purpose and the final constructs generated are listed in Table 1. The required vector was transformed into *Escherichia coli* BL21 (DE3) cells containing the RIL Cam^r^ plasmid (Stratagene) for expression analysis.

**Table 1.**
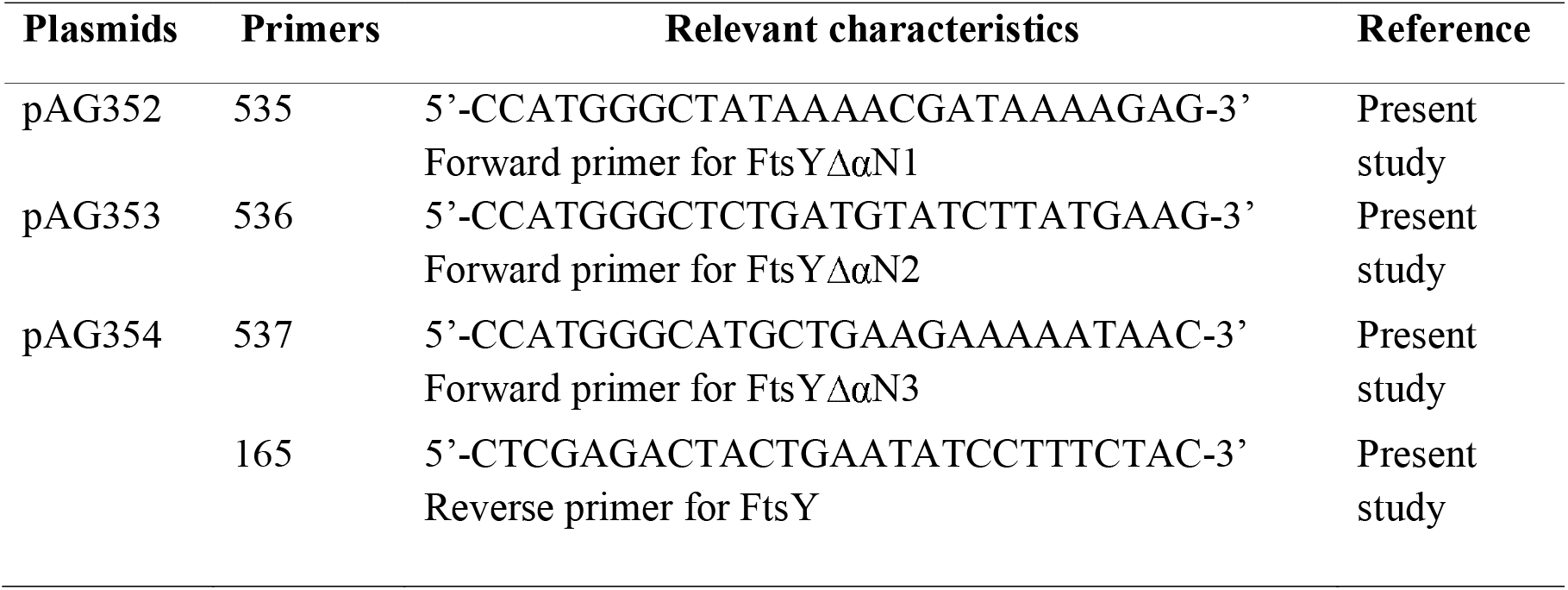
Primers and plasmids used to construct FtsYΔαN mutants.

### 2.2. Expression of recombinant proteins in *E. coli*

Overnight grown culture of *E. coli* BL21(DE3)-RIL cells transformed with the recombinant plasmid were taken in 10 ml volume, and used to inoculate 1 L of Luria–Bertani medium containing kanamycin (50 μg/ml) (for pAG351) or ampicillin (for pAG61-71) and chloramphenicol (34 μg/ml). Cells were grown at 37 °C until OD_600_ reached ~0.6 and then 500 μM IPTG (isopropyl β-D-thiogalactopyranoside) was added. The growth was continued overnight in shaking condition at 16 °C to reduce the inclusion body formation. The cells were collected by centrifugation, resuspended in lysis buffer [50mM Tris-Cl (pH 8.0), 150mM KCl, and 10% glycerol] containing the complete EDTA-free protease inhibitor cocktail (1 tablet/50 ml of lysate; Roche). The lysate was frozen in liquid nitrogen and stored at −80 °C.

### 2.3 Purification of recombinant proteins

The frozen resuspended cell lysates were thawed on ice. Lysozyme (1 mM) was added separately to the lysis buffer. After incubating for 30 min on ice, cells were lysed by sonication with Soniprep150 (DJB Labcare, UK). Cell debris were removed by centrifugation at 15000 rpm for 30 min (rotor SA-300; Sorval RC6+, Thermo Scientific). For the purification purpose, the supernatant was passed through a Ni^2+^-NTA affinity column (Qiagen). The column washed gradually with lysis buffer containing 10 mM and 20 mM imidazole, respectively. The bound protein fraction was eluted in lysis buffer containing 100 mM and 200 mM of imidazole. The eluted fraction was monitored by running reducing SDS-PAGE. The fraction containing the desired protein was dialyzed by buffer exchange using PD columns (Cytiva).

### 2.4. Preparation of archaeosome

Archaeosome from *Sulfolobus acidocaldarius* was prepared by the previously described protocol (Roy et al., 2017). Briefly, *S. acidocaldarius* cells were grown in Brock medium at a pH of 3 and 76°C, and Bligh-Dyer method was used to isolate cell membrane. The extraction solvent was prepared by mixing chloroform with methanol (2:1). A total 3.75 ml of this extraction solvent was then added to 1 ml of cell suspension, followed by rigorous mixing for 10 mins. Further, 1.25 ml of chloroform was added to it followed by rigorous mixing on vortex again, for 1 min. Finally, 1.25 ml of water was added to the mixture and centrifuged at 1000 rpm for 10 mins. The organic layer was collected, and the chloroform was evaporated using a Turbovap LV evaporator (Biotage, Sweden) under nitrogen gas pressure at 65°C. The precipitate was then resuspended in 50 mM phosphate buffer (pH 8.0) preheated at 60°C. The resuspended membrane was snap-frozen in liquid nitrogen followed by thawing in a water bath at 60°C. Archaeosome was prepared from isolated membranes by sonicating the solution for 5 min (10 cycles x 30 sec).

### 2.5. GTPase assay and kinetic analyses

The basal GTPase activity and the kinetics of dual GTP hydrolysis were measured following the protocol discussed earlier (Gupta et al., 2021). Briefly, for the kinetic analyses, 1-10 μM of protein was assayed for the GTPase activity over a range of 5-30 mins. An aliquot of 20 μl from each sample was mixed with 200 μl of Malachite Green reagent and the absorbance was measured at 620 nm spectrophotometrically. The data were corrected for non-enzymatic GTP hydrolysis. The amount of GTP hydrolyzed by individual concentrations of the protein was determined as a function of time, and the resulting data were fitted to the following equation:

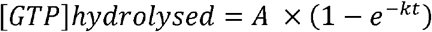

where t is the time (min) of incubation, A is the amount of GTP hydrolyzed at each turnover (μM), and k is the reaction rate, measured in min^-1^.

The affinity of the proteins for GTP can be calculated from the dependence of the observed reaction rate on protein concentration using the following equation:

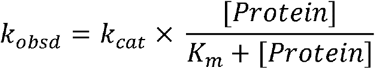

where k_obsd_ is the observed rate constant at a specific protein concentration, k_cat_ is the maximal rate constant with saturating protein, and K_m_ is the protein concentration that provides half the maximal rate.

### 2.6. Tagging of proteins with fluorescent probes

Purified recombinant SRP54 and FtsYΔαN constructs were conjugated with AlexaFluor488 and AlexaFluor532, respectively, following the manufacturer’s protocol (Thermo Scientific). Briefly, SRP54 (50 μM) and FtsY (75 μM) in 50 mM Na-Phosphate buffer supplemented with 100 mM sodium bicarbonate (pH 9.0) were mixed either with AlexaFluor488 or AlexaFluor532 probes. Stock solutions of AlexaFluor dyes were prepared in anhydrous DMSO at 600 μg/ml concentration. For labelling protein samples, 50 μl of fluorescent probe was taken from the stock solution and added very slowly to 1 ml of the protein with gentle and continuous stirring for 2 h at room temperature. Bio-Gel A column was used to separate the labelled protein from the free probe. The fraction, which contained the labelled SRP54 or FtsY, was collected and dialyzed against 50 mM phosphate buffer (pH 7.2) for 24 h at room temperature. The amount of protein labelled with the probe was estimated by measuring the absorbance of the tagged proteins at 280 nm and 495 nm (for AlexaFluor488) or 542 nm (for AlexaFluor532). Finally, tagging of the protein was visualised by running an SDS PAGE followed by scanning in a Typhoon scanner (GE Healthcare, Sweden).

### 2.7. Fluorescence resonance energy transfer (FRET)

To determine the interaction between the two proteins, SRP54 and FtsY, fluorescence resonance energy transfer (FRET) technique was used. The donor AlexaFluor488-labelled SRP54 (100 nM) was incubated for 15 min at 60°C in 50 mM phosphate buffer (pH 7.2) with 100 μM of non-hydrolyzable GTP analog (GppNHp) and 7S RNA (100 nM) and then sequentially titrated with AlexaFluor532-labelled FtsY or its deletion-variants (25-400 nM). The reaction mixture was excited at 495 nm and the fluorescence emission spectra were taken from 505-625 nm at 60°C using a Hitachi f-700 spectrophotometer (Hitachi, Japan). The emission intensity of AlexaFluor488 at 520 nm was measured separately at different titration and was used to calculate the dissociation rate constant (K_d_) from the following equation:

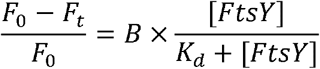

where F_t_ is the fluorescence intensity at 520 nm at different FtsY concentrations, and F_0_ is the fluorescent intensity at [FtsY] = 0; K_d_ is the dissociation rate constant, B is the maximum intensity achieved at saturation of binding.

### 2.8. Membrane fluidity measurement

Membrane fluidity was monitored by the fluorescence anisotropy assay established in the laboratory (Roy et al., 2018). A 10 mM solution of DPH (1,6-Diphenyl-1,3,5-hexatriene) (Sigma) was prepared in DMSO. It was then added to the archaeosome sample with the final DPH concentration being 3 μM. This was further incubated at 16 °C for 10 min. Anisotropy of DPH was then measured in a JASCO fluorimeter (Jasco, Japan) at 35°C. Changes in anisotropy of DPH were monitored with archaeosome mixed either with 10 μM of wildtype and mutant FtsY with or without 10 μM of SRP54. Anisotropy percentage was calculated by considering the anisotropy of the archaeosome as 100% and plotted against each of the additions using Sigma plot 12.0.

### 2.9. Protein secondary structure analysis

Far-UV Circular dichroism spectra of wild-type FtsY and its deletion mutants were collected from 200 to 250 nm using a quartz cell with a 1 mm path length in a CD-spectropolarimeter (JASCO, Japan). Data were collected with a pitch of 1 nm and a scan rate of 50 nm/s. The far-UV spectra were recorded after incubating FtsY or FtsYΔαN1 or FtsYΔαN2 either in presence of archaeosome or SRP54 and archaeosome. The reported spectra for far-UV are the mean of three scans.

### 2.10 Structural modeling

Molecular dynamic simulations have been performed on four *Sulfolobus acidocaldarius* SRP54 bound to FtsY receptor protein complexes; (i) the wild-type variant: full-length FtsY receptor in complex with full-length SRP54, (ii) the N1 variant: Y87-S369 fragment of the FtsY receptor in complex with the full-length SRP54, (iii) the N2 variant: S109-S369 fragment of the FtsY receptor in complex with the full-length SRP54, and (iv) the N3 variant: G129-S369 fragment of the FtsY receptor in complex with the full-length SRP54. Due to the unavailability of experimentally reported structures, ColabFold notebook (Mirdita et al., 2022) has been used to take advantage of the AlphaFold-Multimer model for the building of the initial complexes. The predicted models are in good agreement with the experimentally reported archaeal complex of SRP54-FtsY (PDB: 5L3S). The estimated RMSDs considering the common domains of the built complexes with the archaeal complex are ~1.75 Å for wild type model, ~1.21 Å for the N1 variant, ~1.75 Å for N2 variant, and ~1.9 Å for the N3 variant. In the experimentally reported complex structure, though the FtsY receptor was obtained from *Sulfolobus acidocaldarius,* it was missing the initial M1-L85 fragment. In the AlphoFold predicted wild-type complex, the FtsY unit displayed an RMSD of ~1.44 Å from the experimentally solved FtsY receptor. All the systems were then solvated in a water box of OPC water models (Izadi et al., 2014) where on each side of the protein complex padding of at least ~12 Å of water is maintained. The overall charges of the systems were further neutralized by the addition of an adequate number of Na^+^ ions, and then excess Na^+^ and Cl^-^ ions were added to top up the salt concentration to 0.15 M.

### 2.11. Molecular dynamic simulation

MD simulations were performed using the Amber ff19SB force field parameters (Tian et al., 2020) for the protein molecules. First, all the systems were subjected to minimization to overcome potential inter-and intra-molecular steric clashes. Then, the systems were equilibrated for 200 ps each while imposing positional restraints of 200 kJ/mol. Å^2^ on the protein backbone to relax water molecules and ions around the complexes.

Finally, the backbone restraints were removed and each of the four systems was simulated for 30 ns in 3 replicas each, accumulating ~120 ns of total sampling. All simulations were performed under NPT ensemble by keeping the temperature at 300K using the Langevin thermostat,^15^ with a collision frequency γ = 1 ps^-1^, and pressure at 1 atm using the Monte Carlo barostat (Åqvist et al., 2004). The integration time step of 2 fs has been used after constraining all bonds involving Hydrogen atoms to their equilibrium bond lengths with the SHAKE algorithm (Ryckaert et al., 1977. For all these simulations, short-range nonbonded interactions were calculated using a cutoff at 12 Å, whereas the long-range electrostatic interactions were computed using the Particle Mesh Ewald method (Darden et al., 1993). All the simulations were performed using the *Making it Rain* (Arantes et al., 2021) Google Colab framework that utilizes OpenMM toolkit to run simulations (Eastman et al., 2017).

### 2.12 Calculation of Binding Free Energy

For the comparison of binding free energies between the studied complexes, the molecular mechanics Poisson-Boltzmann surface area (MM/PBSA) method (Kollman et al., 2000) has been employed in each of the simulated ensembles. In this post-processing method, representative snapshots or conformations from an ensemble are used to calculate the free energy change between the bound and free states of a receptor and ligand. Free energy changes of complexation are calculated by combining the gas-phase energy contributions as well as the solvation free energy components (electrostatic and hydrophobic contributions) calculated from an implicit solvent model for each of the units undergoing complexation. Here, we have calculated the free energies from the last 2 ns of each of the replicas simulated in this investigation by selecting snapshots at 20 ps intervals. All analyses were performed with the MMPBSA.py script included in the AmberTools package (Miller et al., 2012). Entropic contributions to free energy have not been estimated in the present study because of the significant computational cost involved in such calculation for large systems like ours. So, the results are only comparable on a qualitative aspect between systems of interest.

## 3. Results

The previously solved crystal structure of the crenarchaeal FtsY (5L3W) provided a composite view of the NG domain heteroxdimer formed by SRP54 and FtsY where the αN1 helix of the receptor was shown to be detached from the core of archaeal TC (Wild et al., 2016). It was found that the extensive electrostatic interaction between the two GTPases is established mostly along the αN2 and αN3 helices (Fig. 1a). Surprisingly, when we modeled the TC in *S. acidocaldarius* without any bound nucleotide, with the help of AlphaFold2, the initial Met1-Gln71 region (excluded in the crystal structure) adopted a helix-turn-helix conformation, which was predicted to include an otherwise disordered acidic domain (Fig. 1b). This raised a vital question regarding the involvement of the acidic domain as well as the αN1 domain in the formation of TC. We, therefore, sought to resolve the issue by investigating the effect of these domains upon SRP54-FtsY interaction. Three different deletion constructs were designed – FtsYΔαN1 (Tyr87-Ser369), FtsYΔαN2 (Ser109-Ser369), and FtsYΔαN3 (Gly129-Ser369) – which were purified using affinity chromatography.

**Figure 1.**
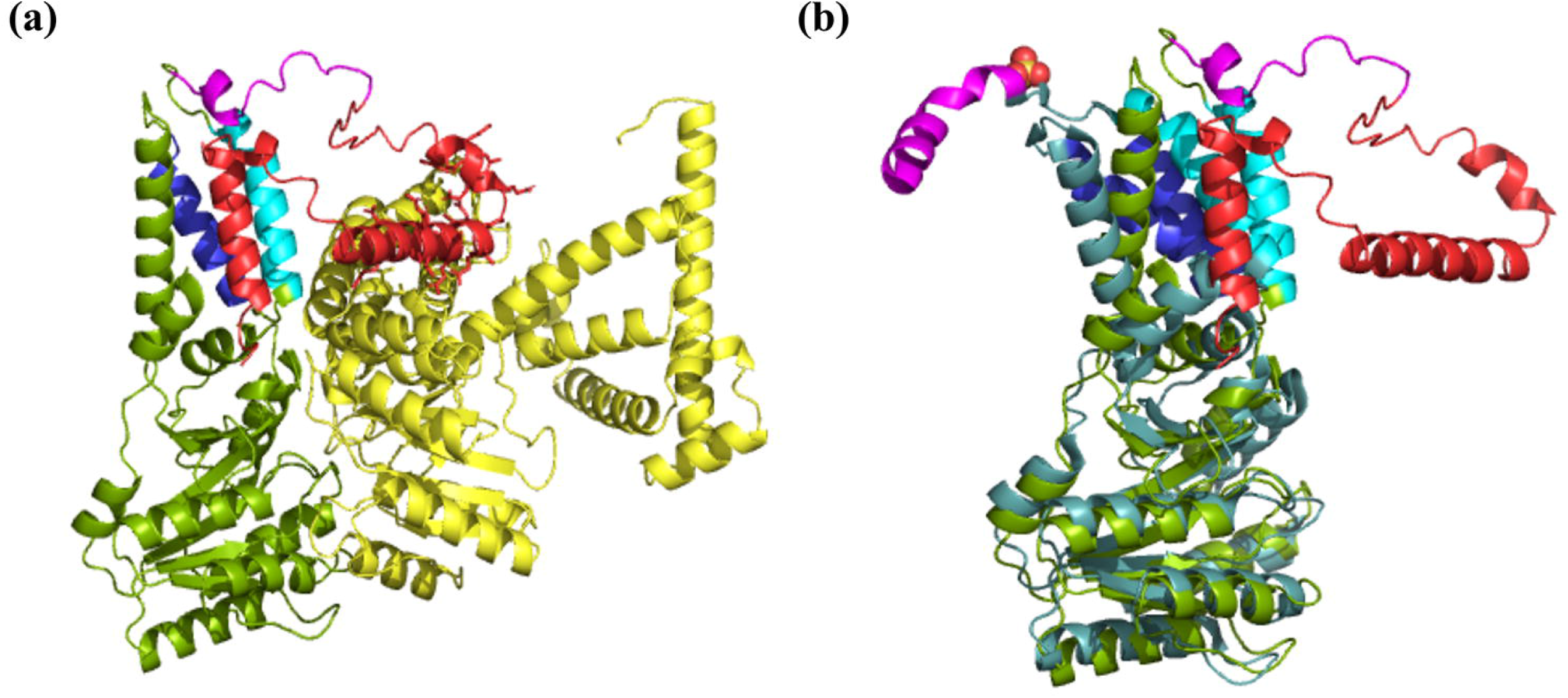
Modeling of archaeal FtsY and purification of its variants. (a) The targeting complex in *Sulfolobus acidocaldarius* was modeled using AlphaFold2 Multimer; SRP54 (yellow) and FtsY (green) are shown in the ribbon diagram. N-terminal motifs are marked in FtsY (A domain, red; αN1, magenta; αN2, cyan; αN3, blue). (b) AlphaFold2 prediction of full length FtsY (colour) aligned with 5L3W (colour) (RMSD = 3.299).

### 3.1. Role of N-terminal alpha-helices in SRP54-FtsY association

Energy transfer due to fluorescence resonance is an important tool that has been employed to assess protein-protein interaction in several assays. The addition of GTP and Ffh was shown to impart an increment and blue shift in the intrinsic tryptophan fluorescence of FtsY (Jagath et al., 2000) which was a direct measure of their association kinetics. Inter-domain interaction, as observed between NG and M domains of Ffh upon FtsY or 4.5S RNA addition, could be followed to saturation using different fluorophore labels (Buskiewicz et al., 2005). The present experiment employed FITC-labelled *Saci*SRP54 and TRITC-labelled variants of FtsY, and it was observed that with each N-terminal alpha-helix deletion, the association between the two proteins was gradually abrogated (Table 2). Since twinning of the nucleotide substrate and SRP RNA binding were found to be decisive in a strong interaction between SRP54/Ffh and FtsY (Egea et al., 2004; Zhang et al., 2008), saturating concentrations of archaeal 7S RNA and non-hydrolyzable GTP analog, GMPPNP, were used in all assays. Unlike GTP, GMPPNP allows capturing the stable targeting complex that would not dissociate following the nucleotide’s hydrolysis. The bacterial and crenarchaeal targeting complexes were previously found to harbor strong electrostatic interactions across the N-terminal alpha-helices in both participating SRP-GTPases (Egea et al., 2004; Ataide et al., 2011; Wild et al., 2016). Deletion of those helices consequently affected the association between the two proteins. The Association of the wildtype TC in *S. acidocaldarius* was recorded previously by a similar FRET experiment which was found to be 40.5 ± 10.8 nM (Gupta et al., 2021). Deletion of 1-72 residues (αN1) augmented the equilibrium dissociation constant for SRP54-binding by almost two folds (K_d_ = 75 ± 5.4 nM). Subsequent deletions of 1-86 residues (αN2) and 1-108 residues (αN3) raised the Kd value to 114.14 ± 24.5 nM and 272.18 ± 47.5 nM, respectively. The highly increased value of the equilibrium dissociation constant possibly indicates that the SRP54-FtsYΔαN3 complex may not be a functional association at all, a consideration that needed to be checked in light of the inherent enzymatic activity of the targeting complex.

**Table 2.**
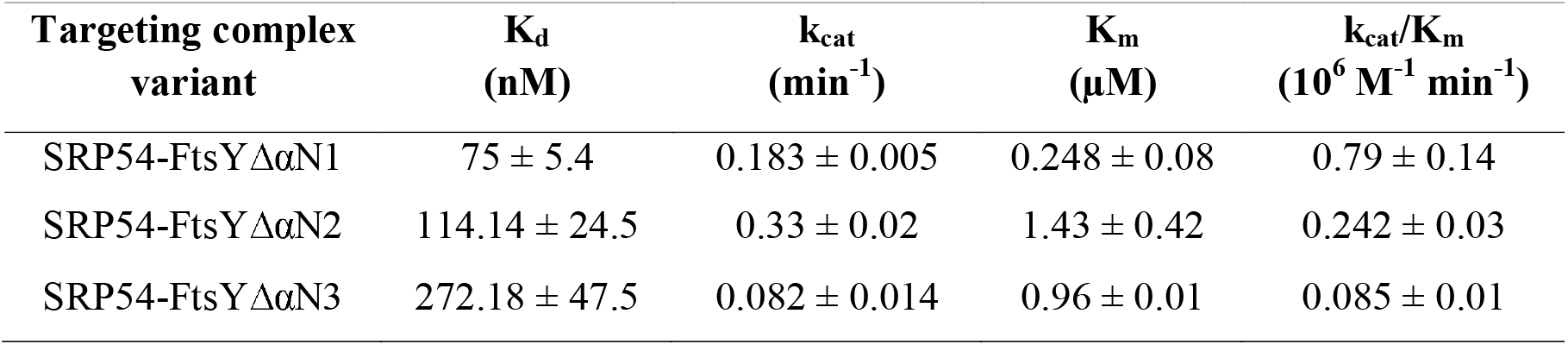
Summary of binding and catalytic constants of TC variants.

**Table 3.**
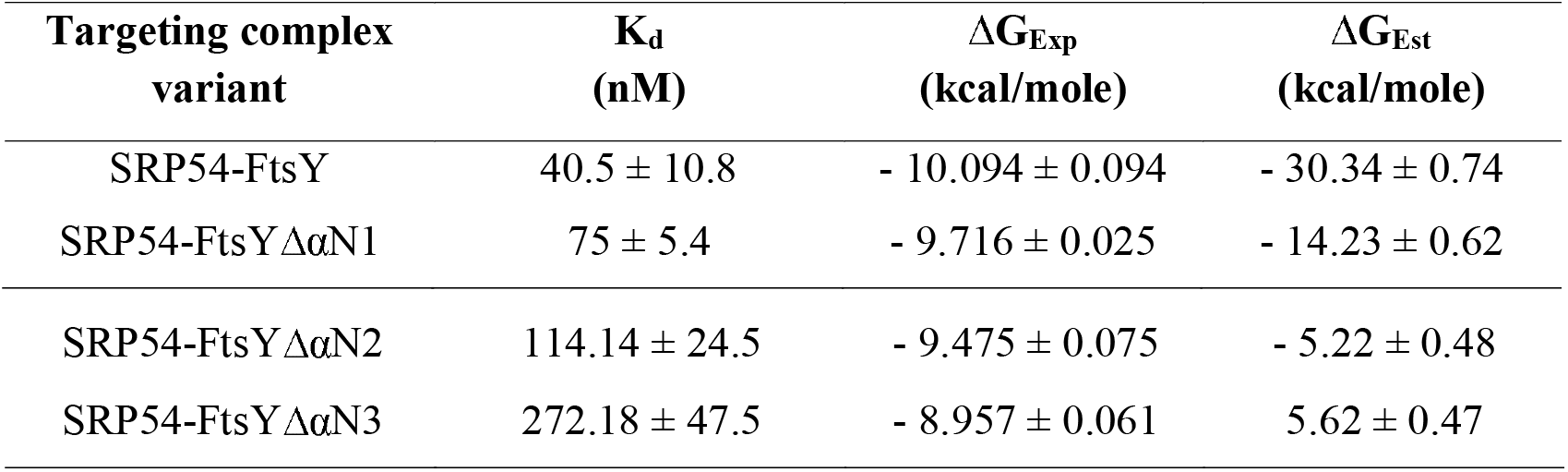
Summary of the experimental and estimated binding free energy.

### 3.2. Reciprocal GTPase activity affected by deletion variants of FtsY

The SRP GTPases are known to reciprocally activate the GTP hydrolyzing ability of each other – an event that is catalyzed by SRP RNA and facilitated by the proper association of the two proteins (Shan et al., 2004; Zhang et al., 2008; Gupta et al., 2021). Several structural studies with both bacterial and archaeal SRP systems have shown that the dual interaction of SRP54-M domain and NG domain with the proximal and distal helices of SRP RNA, respectively, reorient the SRP54 protein in a certain conformation that favors a functional NG-heterodimeric association between SRP54/Ffh and FtsY (Rosendal et al., 2003; Hainzl et al., 2007; Ataide et al., 2011; Voights-Hoffmann et al., 2013). Considering the abrupt changes in the binding phenomenon of the two GTPases because of N-terminal alpha-helix deletions in FtsY, the present experiment sought to characterize the TC variants for their basal GTPase activity. The reactions were set up by the protocol described earlier (Gupta et al., 2021) where a fixed concentration (2.5 μM) of SRP54 was combined with varying concentrations (1-15 μM) of FtsY for assessing the rate of GTP hydrolysis through 0-30 minutes. The resultant rate constants were fit in a modified ligand binding equation and the k_cat_ and K_m_ were calculated accordingly (Table 2). The maximum hydrolytic capacity of the SRP54-FtsYΔαN1 complex was calculated to be 0.183 ± 0.005 min^-1^, which was lower than that of the wildtype SRP54-FtsY complex (0.25 ± 0.01 min^-1^, Gupta et al., 2021). But surprisingly, its affinity for the nucleotide (0.248 ± 0.08 μM), as well as the catalytic efficiency (0.79 ± 0.14 × 10^6^ M^-1^ min^-1^) became a little improved compared to the wildtype complex which has a K_m_ of 0.48 ± 0.12 μM and k_cat_/K_m_ of 0.54 ± 0.07 × 10^6^ M^-1^ min^-1^. The crystal structure of the crenarchaeal FtsY has shown that the αN1 helix is excluded from NG heterodimer in its crystalline state (Wild et al., 2016). This could probably be the reason for the efficient catalytic activity presented by the αN1-deletion variant. The SRP54-FtsYΔαN2 complex showed a lower affinity toward GTP (1.43 ± 0.42 μM) and subsequently a much lower catalytic efficiency (0.242 ± 0.03 × 10^6^ M^-1^ min^-1^) which directly connects to the fact that αN2 helix is directly involved in the interaction between the two GTPases (Wild et al., 2016). The final TC variant, SRP54-FtsYΔαN3, scored the lowest in terms of catalytic activity (0.085 ± 0.01 × 10^6^ M^-1^ min^-1^) though its substrate affinity (0.96 ± 0.01 μM) was comparable to the SRP54-FtsYΔαN2 variant. It was obvious since both αN2 and αN3 helices were deleted in this variant, the overall binding with SRP54 would be largely affected.

### 3.3. Thermodynamic stabilization of targeting complex variants

The consecutive experiments assessing the ability of functional targeting complex formation have projected the αN3-deletion mutant as the least functional FtsY variant. It was evident from the facts that the apparent complexation of this variant with SRP54 was hugely affected in a way that the basal GTPase activity of the complex largely deteriorated. To investigate the probable dynamicity of these TC associations *in silico,* molecular dynamics simulation was carried out. The binding free energy of the protein-protein interaction with Amber-ff19SB was calculated using the standard MM/PBSA technique in three independent runs for each system. The experimental binding energy was calculated as Δ*G* = *RT In K_d_*, where K_d_ is the equilibrium dissociation constant (Table 2). It can be noted that the binding energy predicted by the standard MM/PBSA was much stronger than the experimental energy (Fig. 5), with the value for ΔαN3 variant being positive (ΔG = 5.62 ± 0.47 kcal/mole). This may be attributed to the exclusion of any entropic contribution though, however, studies have shown previously that introducing entropy does not guarantee an obvious improvement in prediction (Sun et al., 2018; Wang et al., 2019).

The N-terminal acidic domain of FtsY (Met1-Leu85) that was excluded in the X-ray diffraction analysis (Wild et al., 2016), took a short alpha-helical confirmation following an initial disordered region in the AlphaFold2 model (Fig. 1a). Consequently, this full-length FtsY variant, complexed with SRP54, showed huge conformational variability in MD simulation which could have contributed to the much stronger binding free energy, ΔG = – 30.34 ± 0.74 kcal mole^-1^, compared to the experimental estimation of −10.094 ± 0.094 kcal mole^-1^. Though the experimental energy estimation for ΔαN1 and ΔαN2 variants (−9.716 ± 0.025 and −9.475 ± 0.075 kcal mole^-1^) lay close to that of the wildtype, their simulated free energy calculations were hugely varying (−14.23 ± 0.62 and −5.22 ± 0.48 kcal mole^-1^). Deletion of all three N-terminal helices has been shown to abrogate the formation of a functional targeting complex and thereafter its activity (Fig. 2, Fig. 3). Thus, positive binding energy possibly hints at the loss of association between SRP54 and FtsYΔαN3 as evident from the much higher K_d_ value of their binding interaction (Table 2). Overall, the simulation experiment suggests that the functional association between SRP54 and FtsY is not permitted beyond the deletion of the αN1 and αN2 helices; also, together the αN1 and the acidic domain may contribute largely to favor the association.

**Figure 2.**
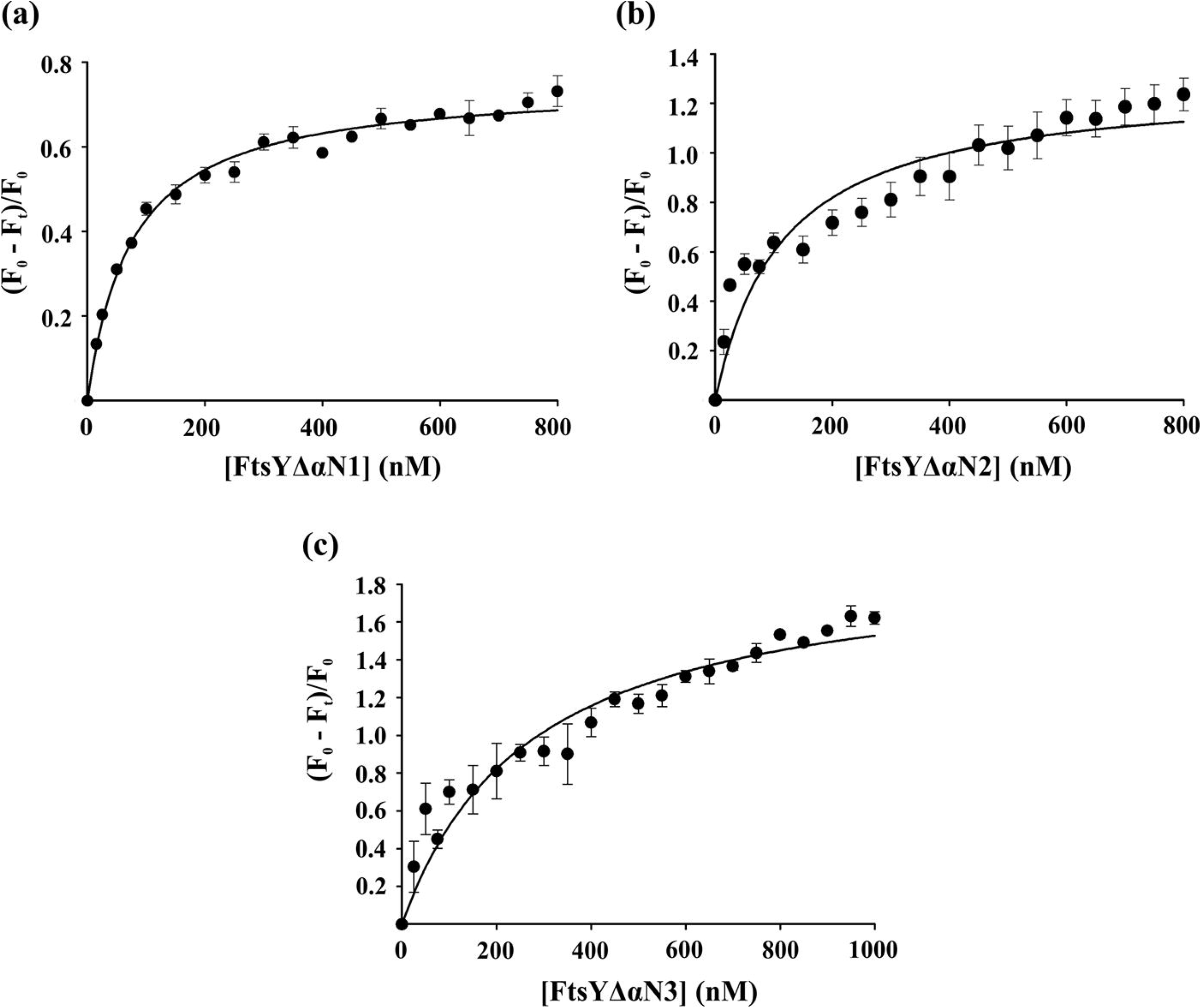
Targeting complex association is governed by N-terminal alpha-helices in FtsY. FRET between FITC-tagged SRP54 (100 nM) and TRITC-tagged FtsY variants (50-400 nM) was carried out in presence of both 100 μM GppNHp and 0.5 μM 7S RNA. The reaction was carried out at λ_Ex_ = 495 nm and data were recorded at λ_Em_ = 520 nm for FtsYΔαN 1 (a), FtsYΔαN2 (b), and FtsYΔαN3 (c).

**Figure 3.**
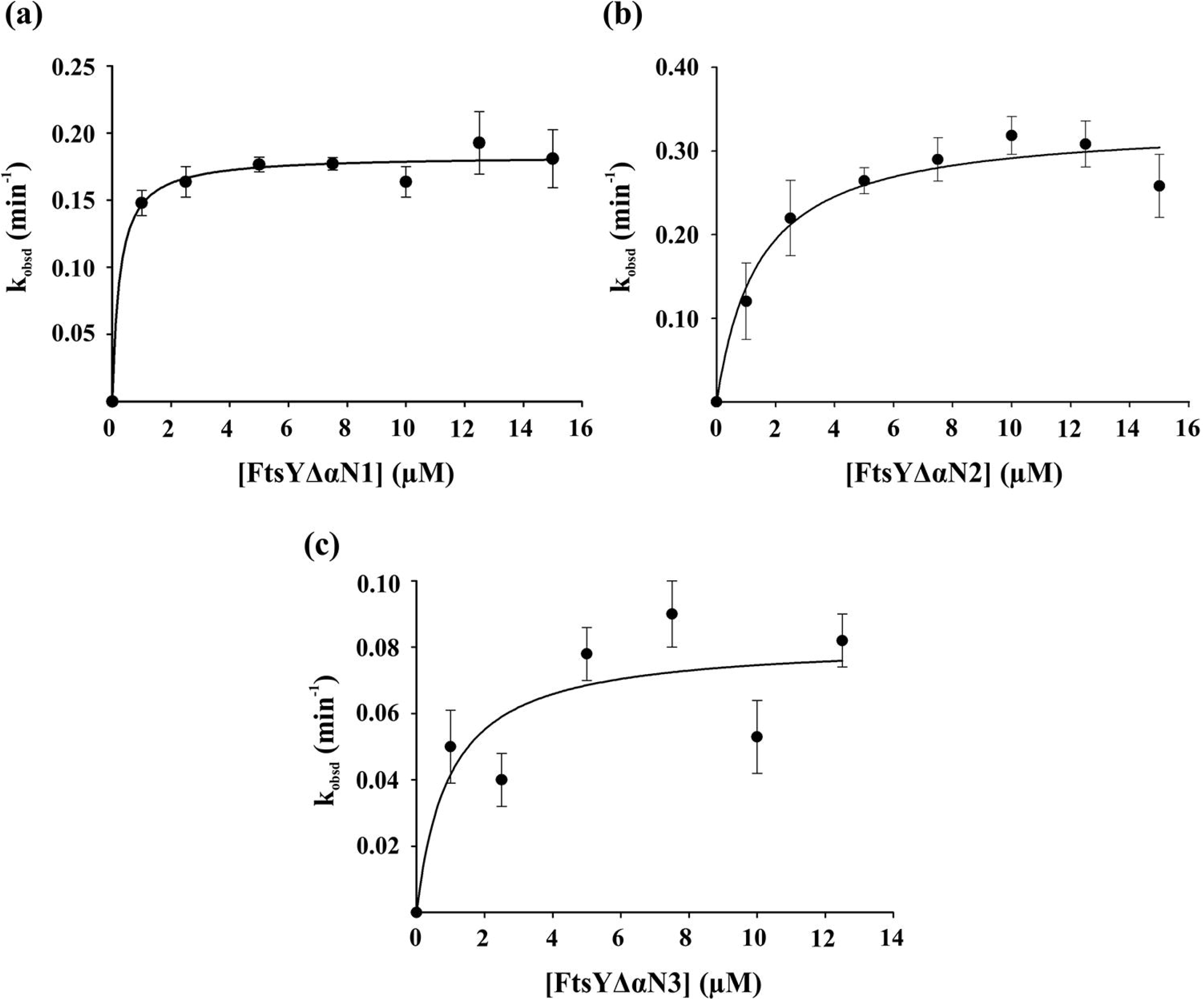
Dual GTPase activity of the targeting complex variants. The reactions were carried out in a series of 0-30 minutes, in the presence of 2.5 μM SRP54. The amount of released phosphate was measured using Malachite-Molybdate reagent and plotted against the time of incubation to generate kobsd. Each reaction was carried out through a fixed concentration range (1-15 μM) for FtsYΔαN1 (a), FtsYΔαN2 (b), and FtsYΔαN3 (c).

### 3.4. Effect of membrane lipid on the conformation of TC variants

Studies in *E. coli* SRP system have shown that the highly charged acidic domain (initial 196 residues) in FtsY is involved in successful membrane targeting (de Leeuw et al., 1997, Zelazny et al., 1997). The otherwise cytosolically distributed bacterial SRP receptor was found to associate strongly with anionic phospholipids, thus proving the importance of negatively charged A domain (de Leeuw et al., 2000; Lam et al., 2010). However, deletion of this domain, but one amino acid, retained most of the functionalities of FtsY in *E. coli* (Stjepanovic et al., 2011; Draycheva et al., 2016). The effect of membrane lipid on the secondary structure of individual peptide segments representing different parts of the membrane targeting sequence (MTS) of bacterial FtsY was assessed by circular dichroism analysis (Stjepanovic et al., 2011) which revealed induction of alpha-helix formation with increasing concentrations of anionic phospholipid. Here, the ΔαN variants of crenarchaeal FtsY were investigated for any change in their apparent secondary structure in presence of archaeosome isolated from *S. acidocaldarius.* The far-UV CD spectra recorded for the wildtype FtsY and both ΔαN variants, in the presence and absence of SRP54 and archaeosome, were presented as a plot of mean residue ellipticity (MRE) versus wavelength (Fig. 6a-c), after subtracting the background signal and the spectrum for SRP54 alone. Based on the previous observations, FtsYΔαN3 was excluded from any further experiment. A decrease in the alpha-helical content of FtsY was evident only after the addition of SRP54 alone or with lipid (Fig. 6a), classically depicted by the decrease in the negative value of MRE at 208 nm and 222 nm (Greenfield NJ, 2007). A similar pattern was observed for FtsYΔαN1 after SRP54 and archaeosome addition but to a lower extent, though the MRE was slightly reduced after the initial addition of archaeosome alone (Fig. 6b). When the MRE at 222 nm (Θ_222_) was plotted against the varying reaction conditions (Fig. 6d), FtsYΔαN1 showed a steady decrease in its alpha-helical content, comparable to the pattern found in the wildtype receptor. The most dramatic change was observed for FtsYΔαN2 as it underwent a sharp increment in its alpha-helical content upon the addition of SRP54. But an incubation with archaeosome and SRP54 reduced the signal to a value similar to the one found in FtsYΔαN2 and FtsY after the addition of archaeosome alone (Fig. 6c-d). Interaction with SRP54 is maintained along the N-terminal alpha-helices, αN2 and αN3, in FtsY. In absence of αN2, αN3 probably attains a more rigid conformation around SRP54 which might result in the apparent increase in the MRE. The exclusion of αN1 from the core NG heterodimer is probably attributing to the slight change in its alpha-helical conformation. FtsY, on the other hand, has the full-length A domain along with the αN1 helix. Structural modeling of the wildtype TC showed the αN1 region to be randomly coiled, as opposed to a helix (Fig. 1a, magenta region). MD simulation also observed a highly dynamic nature of this region in targeting complex formation, which may include the opening of short helices, leading to the observed decrease in MRE for FtsY.

**Figure 4.**
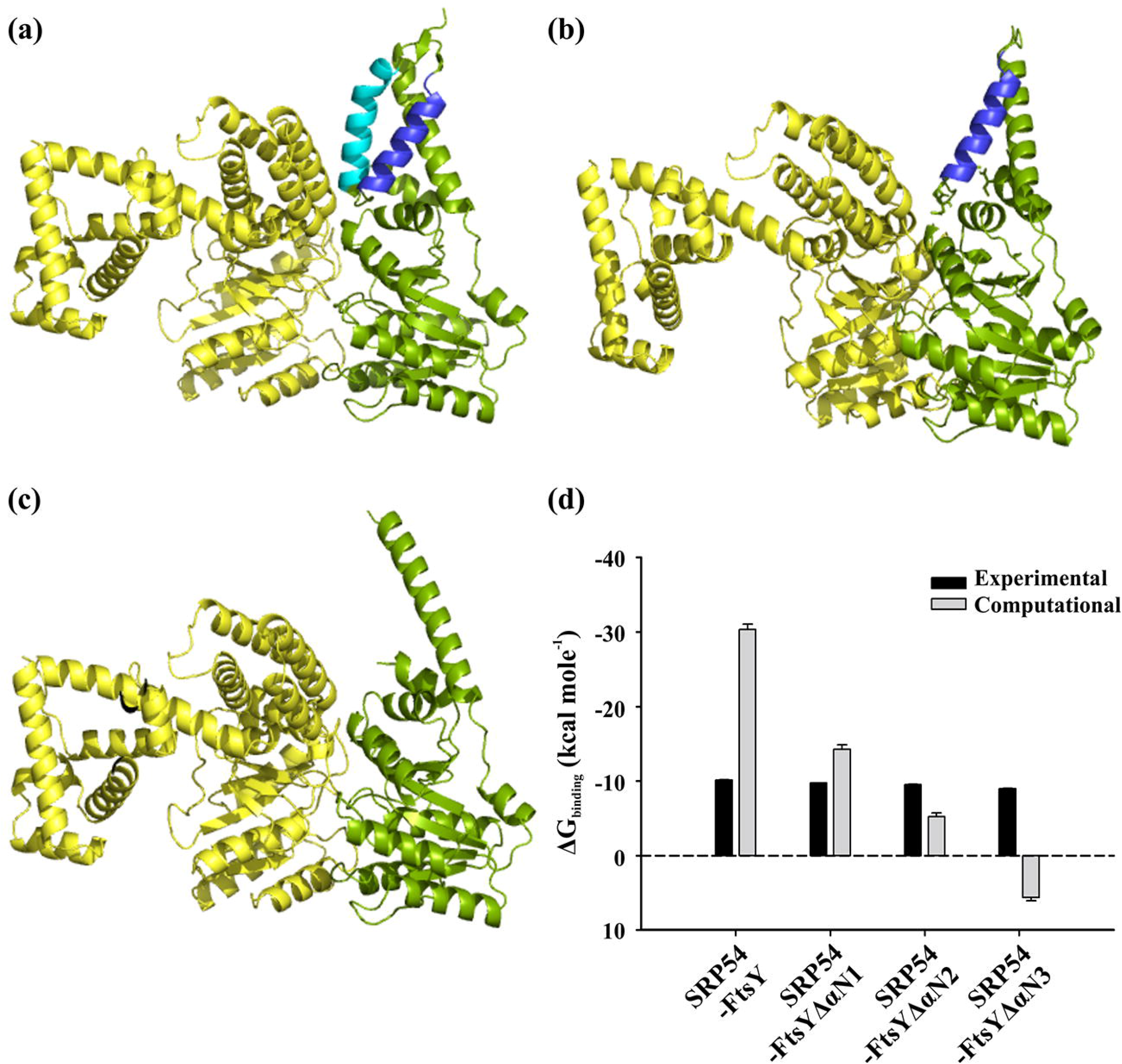
Comparison of the binding profile of targeting complex variants. Each variant was modeled using the AlphaFold2-multimer program and simulated under the Amber-ff19SB force field. The resultant models were processed in PyMol as a complex of SRP54 (yellow) with (a) FtsYΔαN1, (b) FtsYΔαN2, and (c) FtsYΔαN3 (all shown in green). The N-terminal helices are marked as indicated in Fig.1. (d) Overall comparison of the binding free energy of wildtype and mutants obtained from experimental (black) and MM/PBSA calculation (grey).

**Figure 5.**
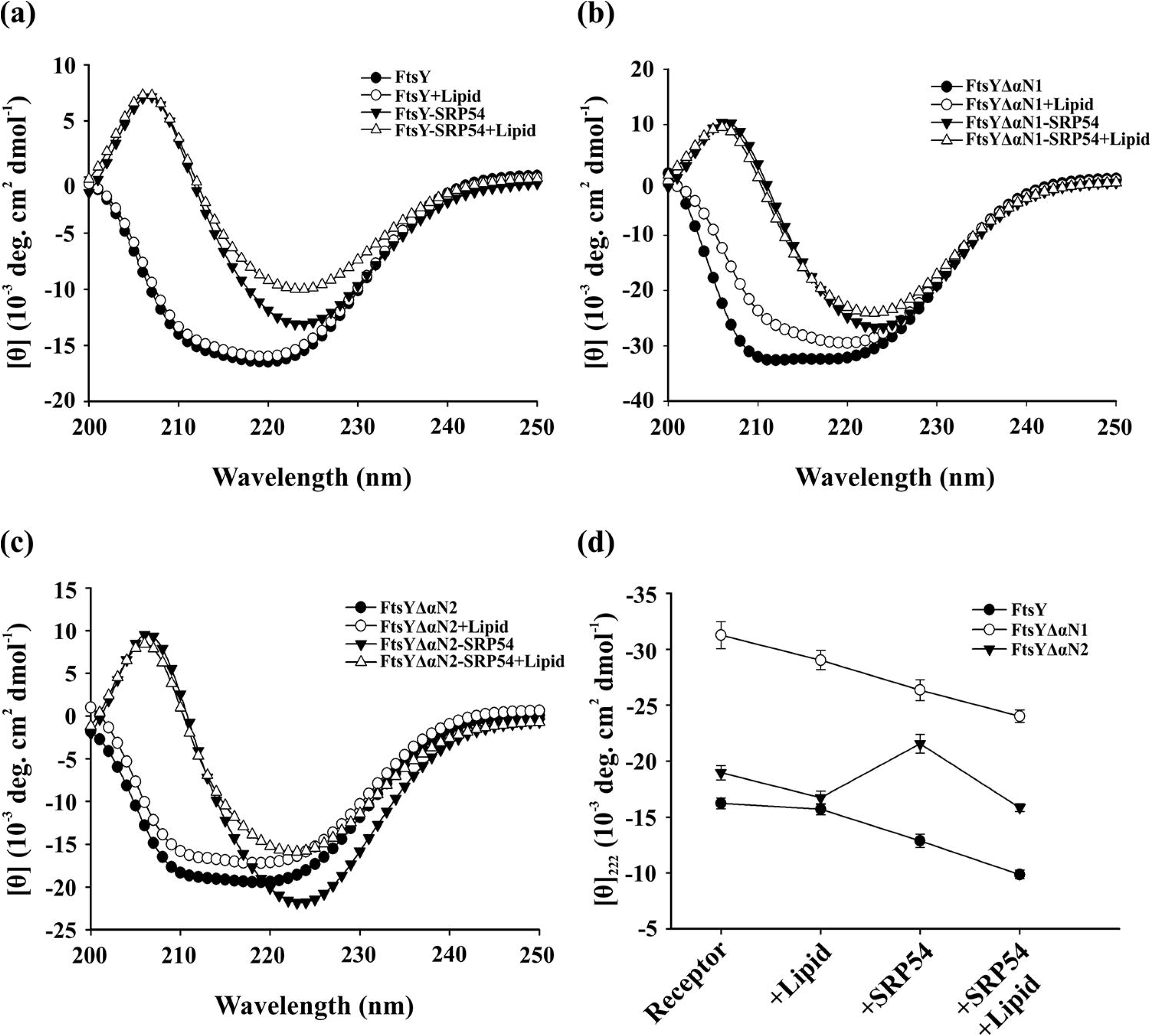
Lipids induce change in helical content of the TC variants. The wildtype, ΔαN1, and ΔαN2 variants of FtsY were analyzed in presence of archaeosome, SRP54, and both SRP54 and archaeosome. The spectrum was recorded at 200-250 nm range and 35 °C for FtsY (a), FtsYΔαN1 (b), and FtsYΔαN2 (c) and represented as mean residue ellipticity. (d) The overall transition of the secondary structure for the variants was shown as a function of the reaction conditions.

**Figure 6.**
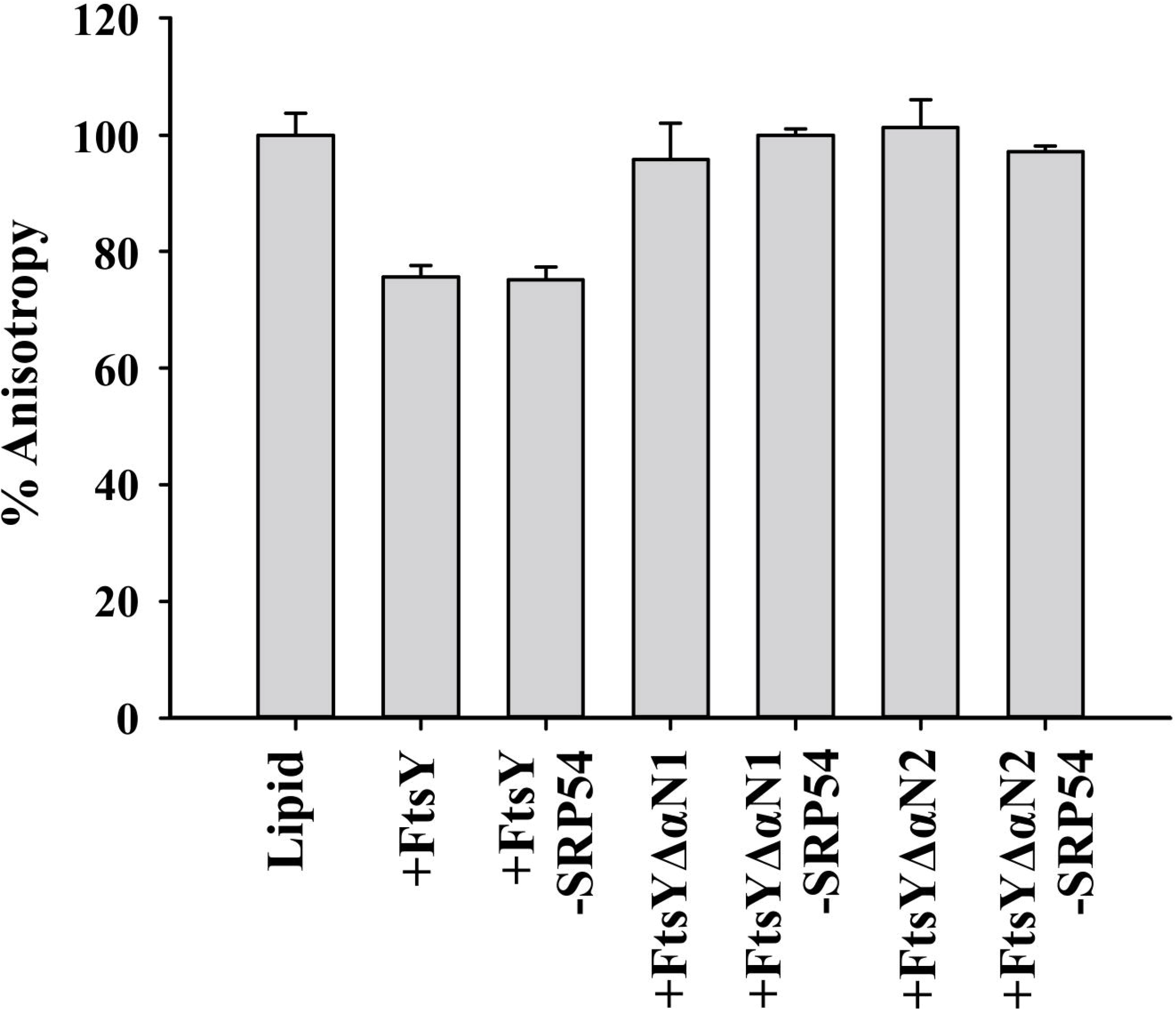
Membrane fluidity is affected by functional targeting complex. Changes in the membrane fluidity of isolated archaeosome fraction upon addition of different protein assemblies were evaluated by measuring the alteration in DPH anisotropy. Individual measurements were taken for FtsY, FtsYΔαN1, FtsYΔαN2, SRP54-FtsY, SRP54-FtsYΔαN1 and SRP54-FtsYΔαN2 at λ_Ex_ = 350 nm and λ_Em_ = 430 nm.

### 3.5. SRP receptor influences membrane fluidity

A close correlation has been observed between the lipid association of membrane targeting proteins and membrane fluidity (Kremer et al., 2001; Roy et al., 2018). To determine if FtsY interacts with membrane phospholipids via an integral domain, we tested the effect of the receptor and its ΔαN variants on archaeosome formed of lipids extracted from *S. acidocaldarius* cells. To assess membrane fluidity, the lipophilic probe DPH was tagged to the archaeosome preparation. The steady-state fluorescence anisotropy of this probe is inversely related to the fluidity of the membrane and is thus used to monitor the dynamic properties of the membrane. Since DPH attaches to the hydrophobic core of the lipid bilayer, its motional order within the core affects its inherent fluorescence property. The current experiment presents a comparative account of the anisotropy measurement taken in presence of different variants of the SRP receptor and the targeting complex (Fig. 7). The anisotropy of the labeled archaeosome (R = 0.0598 ± 0.003) was taken as 100% and the rest of the values were normalized accordingly. It showed that the addition of FtsY and SRP54-FtsY complex decreased the anisotropy value by ~25%, whereas any of the ΔαN variants, alone or in complex with SRP54, did not alter the DPH anisotropy to any significant extent. The reduced anisotropy values were measures of increased fluidity of the bilayer in presence of the wildtype receptor and targeting complex, proving that initial N-terminal domains of the A-αN1 region may be critical for membrane association.

## 4. Discussion

The signal recognition particle system across all domains of life employs two special GTPases – SRP54/Ffh and SR/FtsY. Bypassing the need for an external guanine nucleotide exchange factor, these two proteins make use of the unique insertion box domain present in their classical GTPase motifs, which acts as an internal tool to control the exchange of GDP and GTP (Moser et al., 1997), and reciprocally activate each other (Egea et al., 2004; Focia et al., 2004). For a successful turnover of this reciprocal activation, the two GTPases need to associate together to form a functional targeting complex with a composite catalytic core formed by electrostatic interaction between their respective NG domains (Gupta et al., 2016). Studies with crystal structures of SRP receptors in bacterial and archaeal domains have identified the N-terminal alpha-helices in the NG domains of both proteins to be the key players in mediating this extensive interaction (Ataide et al., 2011; Wild et al., 2016), though concrete biochemical evidence is absent in archaeal SRP system. Apart from the C-terminal NG domain, FtsY has a highly acidic N-terminal A domain that has always been hypothesized to be randomly disordered in solution. Although the importance of this domain in targeting complex association has not been clear, with some groups reporting proper cellular functioning in absence of A domain (Haddad et al., 2005) while some others show abrogated SRP activity (Zelazny et al., 1997), its role in membrane targeting has been thoroughly established in bacteria (Lam et al., 2010; Draycheva et al., 2018).

The present work sought to establish the biochemical basis of the structural contribution of the FtsY-N domain motifs in the functional association of the crenarchaeal targeting complex. The three-terminal helices, αN1, αN2, and αN3, were sequentially deleted to construct three subsequent deletion variants – FtsYΔαN1, FtsYΔαN2, and FtsYΔαN3. These three variants were tested for their ability to bind SRP54 and catalyze dual GTP hydrolysis in complex and their probable association with archaeal membrane lipids was also checked to identify the probable membrane-targeting domain in FtsY. The formation of NG heterodimer between SRP and its receptor is thought to reorient the G domains to form a composite active site at the dimer interface (Shan & Walter, 2005; Egea et al., 2008) that facilitates the reciprocal GTP hydrolysis. Wild et al. (2016) have shown that the αN1 helix of the crenarchaeal FtsY is excluded from the targeting complex association in its crystalline state and the maximum interaction is intensified in the αN2 and αN3 helices. To establish our system to be in total agreement with the already published data, we solved the solution structure of FtsY by the small-angle X-ray scattering analysis. SAXS data were fit into the corresponding GNOM plot and the resulting χ^2^ value of 1.24 indicated a very well match of the solution and crystal structure data. The binding of SRP54 with different ΔαN variants was assessed using the FRET tool and the maximally affected variant was found to be FtsYΔαN3. The apparent K_d_ values of the other two variants indicated that SRP54-binding was moderately compromised for FtsYΔαN2, whereas the ΔαN1 variant bound SRP54 like SRP54-FtsY binding in absence of RNA. The combinatorial GTPase activity of the SRP54-FtsY complex and its ΔαN variants further showed that the SRP54-FtsYΔαN3 complex was highly nonfunctional (k_cat_/K_m_ = 0.085 ± 0.01 × 10^6^ M^-1^ min^-1^) in terms of GTP hydrolysis, whereas the SRP54-FtsYΔαN1 complex showed a slightly higher catalytic efficiency (0.79 ± 0.14 × 10^6^ M^-1^ min^-1^) as compared to the wildtype TC. Since αN1 helix was shown to be excluded from the functional association of crenarchaeal targeting complex (Wild et al., 2016), deletion of this specific segment probably could not affect the functionality of the targeting complex. The evident dynamic nature of this association led us to investigate the thermodynamic behavior of these complexes *in silico* by MD simulation using Open MM protocol. The resultant binding free energy values and the dynamic model corresponding to each complex were surprisingly indicative of a huge conformational rearrangement around the αN1 helix and the A domain preceding it. The SRP54-FtsY complex modeled by the AlphaFold2 Multimer program had the N-terminal 1-85 residues arranged in a helix-turn-helix motif, which was excluded from the previously solved crystal structure (5L3W, 5L3S) and proposed to be disordered in the A domain (1-71 residues) region. The simulated model showed very high conformational variations in the wildtype TC model, thus leading to a higher value of binding free energy (ΔG). SRP54-FtsYΔαN3 showed a positive value for ΔG, drastically opposite to the experimental finding. This may be due to the severely affected binding interaction or the exclusion of interaction entropy from the calculation. However, deletion of all the αN helices together clearly seemed detrimental to the functional TC formation.

Bacterial FtsY has been studied extensively to pin the translocon binding and membrane targeting function on its A domain (de Leeuw et al., 2000; Draycheva et al., 2016) and to establish the minimal targeting sequence necessary for that function (Lam et al., 2010; Fu et al., 2017). Bacterial FtsY has been shown to prefer anionic phospholipids which tend to aggregate in presence of the receptor (de Leeuw et al., 2000). Although membrane interaction could be achieved by both A and NG domains (Draycheva et al., 2016), the dynamics of the interaction may be different. The native conformation of the A domain was found to be unfolded in solution which undergoes a random coil-helix transition in presence of anionic phospholipids (Stjepanovic et al., 2011) and this interaction is further strengthened when the receptor attains a stable complex with SRP-RNC (Lam et al., 2010; Fu et al., 2017), mostly mediated by the αN1 helix. In the present work, we sought to follow the changes, if any, incurred by the archaeal membrane lipid upon the different TC variants constructed on purpose. To identify changes in the secondary structure of the protein far-UV CD spectrum was obtained, with or without archaeosome, for receptor variants alone and in complex with SRP54. The alpha-helical contents of the wildtype and mutant TCs were found to decrease in presence of lipids, with the maximum effect in the wildtype complex with full-length FtsY. This observation clearly showed that the unfolding of the initial A-αN1 region is triggered by archaeosome, contrary to the findings in bacterial FtsY. A similar observation could be obtained from the membrane stabilization assay where the fluidity of the lipid bilayer is measured by changes in steady-state fluorescence anisotropy of a bilayer-binding probe, DPH. A ~25% reduction in anisotropy only in presence of wildtype SRP receptor and receptor-SRP54 complex hinted at the possible involvement of the A-αN1 region in membrane interaction.

## 5. Conclusion

The present work sought to identify the minimal domain of FtsY required to associate with SRP54 and archaeal cell membrane functionally. Deletion of the N-terminal alpha-helices resulted in truncated FtsY variants that bound to SRP54 with differing capacities. Also, the reciprocal GTPase activity by the complexes comprising SRP54 and FtsYΔαN variants was severely affected. The pattern of these observations suggested that deletion of all three helices is detrimental to native functions of FtsY, but deletion of the αN1 helix could retain considerable activities. The binding free energy of the wildtype and mutant targeting complexes were computationally calculated by MD simulation and compared with the experimental values, which further established the functionality of αN1 helix and the region preceding it (A domain) in the stabilization of the targeting complex. Finally, the mutual influence of archaeal membrane and targeting complexes was analyzed by far-UV CD and DPH anisotropy. Both techniques successfully established that lipid interaction in archaea employs the N-terminal A-αN1 region of FtsY which probably gets relaxed of its helical constraints while associating with the membrane.

